# Urocortin 3 in the posterodorsal medial amygdala mediates psychosocial stress-induced suppression of LH pulsatility in female mice

**DOI:** 10.1101/2021.06.20.449139

**Authors:** Deyana Ivanova, Xiao-Feng Li, Caitlin McIntyre, Yali Liu, Lingsi Kong, Kevin T O’Byrne

## Abstract

Exposure to psychosocial stress disrupts reproductive function and interferes with pulsatile luteinising hormone (LH) secretion in mammals. The posterodorsal sub-nucleus of the medial amygdala (MePD) is part of the limbic brain and is an upstream modulator of the reproductive axis as well as stress and anxiety states. Corticotropin releasing factor type-2 receptors (CRFR2) are activated in the presence of psychosocial stress together with an increased expression of the CRFR2 ligand Urocortin3 (Ucn3) in MePD of rodents. We investigate whether Ucn3 signalling in the MePD is involved in mediating the suppressive effect of psychosocial stress exposure on LH pulsatility. Firstly, we administered Ucn3 into the MePD and monitored the effect on pulsatile LH secretion in ovariectomised mice. Next, we delivered Astressin2B, a highly selective CRFR2 antagonist, intra-MePD in the presence of predator odor, 2,4,5-Trimethylthiazole (TMT) and examined the effect on LH pulses. Subsequently, we virally infected ovariectomised Ucn3-cre-tdTomato mice with inhibitory DREADDs targeting the MePD Ucn3 neurons while exposing the mice to TMT or restraint stress and examined the effect on LH pulsatility as well as corticosterone (CORT) release. Administration of Ucn3 into the MePD dose-dependently inhibited pulsatile LH secretion and intra-MePD administration of Astressin2B blocked the suppressive effect TMT on LH pulsatility. Additionally, DREADDs inhibition of MePD Ucn3 neurons blocked TMT and restraint stress-induced inhibition of LH pulses as well as CORT release in the presence of TMT. These results demonstrate for the first time that Ucn3 neurons in the MePD mediate psychosocial stress-induced suppression of the GnRH pulse generator and psychosocial stress-induced CORT secretion. Ucn3 signalling in the MePD plays a fundamental role in modulating the hypothalamic-pituitary-ganadal and hypothalamic-pituitary-adrenal axes, and this brain locus may represent a nodal centre in the crosstalk between the reproductive and stress axes.

## Introduction

Psychological stress is known to have deleterious effects on reproductive function in mammals, including humans (1). Rodent models of psychosocial stress, such as restraint and predator odor exposure, show reduced luteinizing hormone (LH) pulse frequency (2,3) and LH surge release (4) as well as delayed puberty (5). Kisspeptin (kiss1) neurons located in the arcuate nucleus (ARC) of the hypothalamus are a crucial component of the gonadotropin-releasing hormone (GnRH) pulse generator regulating pulsatile secretion of GnRH to maintain function of the hypothalamic pituitary gonadal (HPG) axis (6–8). These neurons are known as KNDy because they co-express neurokinin B (NKB) and dynorphin A (DYN) and their synchronous activity release pulses of kisspeptin that act on GnRH dendrons in the median eminence to release pulses of GnRH release (9). It has been shown that acute restraint stress rapidly suppresses LH pulses and reduces c-fos in KNDy neurons in mice (10). The underlying neural mechanisms by which psychological stress disrupts the GnRH pulse generator remains to be fully elucidated.

The amygdala, a key part of the limbic brain involved in emotional processing is known to modulate the HPG axis, including exerting an inhibitory break on puberty, where lesioning of the amygdala leads to advancement of puberty in female rhesus macaques (11). Moreover, specific lesioning the posterodorsal sub-nucleus of the medial amygdala (MePD) advances puberty in the rat (12). The majority of efferent projections from the MePD are GABAergic and a significant population project to reproductive neural centres in the hypothalamus (13–16). This amygdaloid nucleus is also known to project to the KNDy neural population in the ARC (17,18). Extrahypothalamic kiss1 neuronal populations exist in the MePD and we have recently shown that optogenetic stimulation of these neurons increases LH pulse frequency in mice (19).

The amygdala is also implicated in the neuroendocrine response to stress, activating the hypothalamic pituitary adrenal (HPA) axis and modulating anxiety behaviour. Restraint stress has been seen to increase c-fos expression in the medial amygdala (MeA) (20), and lesioning this nucleus attenuates restraint-induced suppression of pulsatile LH secretion in rats (2). Moreover, the amygdala processes olfactory information and is robustly activated by predator odor (5,21,22). Particularly, the MePD exhibits a distinct firing pattern in the presence of predator odor and multiunit recordings reveal the MePD is activated in response to predator urine exposure (23). Recently, exposure to cat odor was shown to induce a fear response delaying puberty in rats (5). Therefore the MePD may be involved in the integration of anxiogenic signals with the GnRH pulse generator.

Urocortin 3 (Ucn3), part of the corticotropin-releasing factor (CRF) superfamily, is a highly specific endogenous ligand for CRF type 2 receptors (CRFR2) (24). A high density of Ucn3 and CRFR2 expressing neurons are found in the MePD (25–27). The MePD is part of the mammalian social brain network and selective optogenetic activation of Ucn3 neurons in this region influences social behavior (27). Ucn3 is involved in modulating the stress response and exposure to restraint stress increases MePD Ucn3 mRNA expression in rodents (28). Social defeat, a classical rodent psychosocial stress model, increases CRFR2 expression in the MePD (29).

In this study, we aimed to determine whether Ucn3/CRFR2 signalling in the MePD mediates the inhibitory effect of predator odor and restraint stress on pulsatile LH secretion in adult ovariectomised mice. Initially, we determined the effect of intra-MePD administration of Ucn3 *per se* on pulsatile LH secretion. We then investigated whether intra-MePD administration of the highly selective CRFR2 antagonist, Astressin2B, would block predator odor stress-induced suppression of pulsatile LH secretion. Finally, we employed pharmacosynthetic DREADDs (designer receptor exclusively activated by designer drugs) to selectively inhibit Ucn3 neurons in the MePD of Ucn3-Cre-tdTomato mice aiming to investigate whether endogenous MePD Ucn3 signalling mediates predator odor or restraint stress-induced suppression of LH pulsatility.

## Methods

### Mice

Female C57Bl6/J mice were purchased from Charles River Laboratories International, Inc. Cryopreserved sperm of Ucn3-cre mice (strain Tg(Ucn3-cre)KF43Gsat/Mmucd; congenic on C57BL/6 background) was acquired from MMRRC GENSAT. Heterozygous transgenic breeding pairs of Ucn3-Cre-tdTomato mice were recovered via insemination of female C57Bl6/J mice at King’s College London. Ucn3-Cre mice were genotyped using PCR for the detection of heterozygosity (primers 5’ - 3’: Ucn3F – CGAAGTCCCTCTCACACCTGGTT; CreR – CGGCAAACGGACAGAAGCATT). Ucn3-cre-tdTomato reporter mice were generated by breeding Ucn3-cre mice with td-Tomato mice (strain B6.Cg-Gt(ROSA)26Sortm9(CAG-tdTomato)Hze/J; congenic on C57BL/6 background) acquired from The Jackson Laboratory, Bar Harbor, ME, USA. The reporter allele encoding tdTomato is expressed upon Cre-mediated recombination to obtain heterozygous Ucn3-cre-tdTomato mice, as previously described (27) (primers 5’ - 3’: TA Wild type Forward oIMR9020 - AAGGGAGCTGCAGTGGAG; TC Wild type Reverse oIMR9021 - CCGAAAATCTGTGGGAAG; Mutant Reverse WPRE oIMR9103 - GGCATTAAAGCAGCGTATCC; Mutant Forward tdTomato oIMR9105 - CTGTTCCTGTACGGCATGG). Female mice aged between 6-8 weeks and weighing between 19-23 g were singly housed in individually ventilated cages sealed with a HEPA-filter at 25 ± 1 °C in a 12:12 h light/dark cycle, lights on at 07:00 h. The cages were equipped with wood-chip bedding and nesting material, and food and water ad libitum. All procedures were carried out following the United Kingdom Home Office Regulations and approved by the Animal Welfare and Ethical Review Body Committee at King’s College London.

### Stereotaxic adeno-associated-virus (AAV) injection and cannula implantation

All surgical procedures were carried out under general anaesthesia using ketamine (Vetalar, 100 mg/kg, i.p.; Pfizer, Sandwich, UK) and xylazine (Rompun, 10 mg/kg, i.p.; Bayer, Leverkusen, Germany) and with aseptic conditions. Mice were secured in a David Kopf stereotaxic frame (Kopf Instruments, Model 900) and bilaterally ovariectomised (OVX). To assess the effects of Ucn3 and Astressin2B (Ast2B) in the MePD unilateral and bilateral cannula implantation, respectively, was performed using robot stereotaxic system (Neurostar, Tubingen, Germany). The mouse brain atlas of Paxinos and Franklin (30) was used to obtain target coordinates for the MePD (2.30 mm lateral, −1.55 mm from bregma, at a depth of −4.94 mm below the skull surface). To reveal the skull a midline incision in the scalp was made, the position of MePD was located and a small hole was drilled in the skull above it. A unilateral or bilateral 26-gauge guide cannula (Plastics One, Roanoke, VA, USA) was targeted to the MePD. Once in position the guide cannula was secured on the skull using dental cement (Super-Bond Universal Kit, Prestige Dental, UK) and the incision of the skin was closed with suture. Ten mice were implanted with a unilateral cannula and 14 mice were implanted with a bilateral cannula. Following a one-week post-surgery recovery period, mice were handled daily to acclimatise to experimental procedures.

Intra-MePD bilateral stereotaxic viral injection of the inhibitory adeno-associated-virus carrying the DIO-hM4D, DREADD, construct (AAV-hSyn-DIO-HA-hM4D(Gi)-IRES-mCitrine, 3×1011 GC/ml, Serotype:5; Addgene) was performed for the targeted expression of hM4D-mCitrine in MePD Ucn3 neurons. Initially, Ucn3-Cre-tdTomato mice were secured in a Kopf stereotaxic frame (Kopf Instruments) and bilaterally OVX. The skull was revealed and two small holes were drilled above the location of the MePD. The same coordinates were used to target the MePD, as described above. AAV-hSyn-DIO-hM4D(Gi)-mCitrine (300 nL) was bilaterally injected, over 10 min, into the MePD using a 2-μL Hamilton micro syringe (Esslab, Essex, UK). The needle was left in position for a further 5 min and lifted slowly over 1 min. Cre-positive mice received the AAV-hM4D injection (test mice) or a control virus AAV-YFP (Addgene) (control mice). The control virus does not contain the DIO-hM4D construct. The mice were left to recover for one week. Ten mice received the AAV-hM4D injection and 5 mice received the control AAV-YFP. Following the one-week recovery period, mice were handled daily to acclimatise to experimental procedures for a further 2 weeks.

### Intra-MePD Ucn3 administration and blood sampling for measurement of LH

In order to test the effect of Ucn3 (Cambridge Bioscience, Cambridge, UK) administration into the MePD on LH pulsatility, OVX mice implanted with a unilateral cannula on the right side were subjected to the tail-tip blood collection procedure, as previously described (31). Infusion of drugs and blood sampling were performed between 09:00-13:00 h, where 5 μl blood was collected every 5 min for 2 h 30 min. Unilateral internal cannula (Plastics One) attached to extension tubing (0.58 mm ID, 0.96 mm OD), preloaded with Ucn3 or aCSF as vehicle control, was inserted into the guide cannula. The internal cannula extends 0.5 mm beyond the tip of the guide cannula to reach the MePD. The tubing extended beyond the cage and the distal ends were attached to 10-μl Hamilton syringes (Waters Ltd, Elstress, UK) fitted into a PHD 2000 Programmable syringe pump (Harvard Apparatus, Massachusetts, US), allowing for a continuous infusion of the drug at a constant rate. The mice were kept in the cage throughout the experiment, freely-moving with food and water. After a 60 min control blood sampling period, the mice were given an initial bolus injection of Ucn3 at 112.20 fmol or 1.12 pmol in 0.60 μl at a rate of 0.12 μl/min over 5 min, followed by a continuous infusion of 224.40 fmol or 2.24 pmol in 1.20 μl of Ucn3 delivered at a rate of 0.04 μl/min for 30 min. Ucn3 infusions were performed between 60-95 min of the total blood sampling period. Administration of aCSF was performed in the same manner as a control. The study was cross-designed so that mice receive both doses of Ucn3 and aCSF in a random order, with at least 2 days between experiments.

### Ast2B delivery into the MePD during predator odor-exposure and blood sampling for measurement of LH

To test the effect of acute predator odor exposure on LH pulsatility, OVX mice implanted with a bilateral cannula were exposed to 2,4,5-Trimethylthiazole (TMT; synthetic extract of fox urine; Sigma-Aldrich, UK) while blood samples were collected every 5 min for 2 h, as previously described. Experiments were performed between 10:00-13:00 h. After 1 h of controlled blood sampling, 12 μl of TMT (≥98% purity) was pipetted on a small circular piece of filter paper and placed in a petri dish in the centre of the cage, with blood sampling continued for the remainder of the experiment.

In order to test whether Ast2B (Tocris Bio-techne, Abingdon, UK) administration into the MePD can block the TMT effect on LH pulsatility, OVX mice implanted with a bilateral cannula were acutely exposed to TMT as described above while Ast2B was delivered into the MePD (as described for Ucn3 infusion above) and blood samples were collected. After a 50 min control blood sampling period, mice were given an initial bolus dose of 8.00 pmol Ast2B in 0.50 μl delivered at a rate of 0.10 μl/min over 5 min, followed by a continuous infusion of 19.20 pmol Ast2B in 1.20 μl at a rate of 0.05 μl/min for the remaining 65 min of experimentation. At 60 min, mice were exposed to TMT for the remaining 1 h duration of the experiment, as previously described. The administration of Ast2B only was performed in the same manner excluding TMT exposure, as control. This study was cross-designed, as described above.

### DREADD inhibition of Ucn3 neurons in the MePD in the presence of TMT or restraint stress and blood sampling for measurement of LH and CORT

We used OVX Ucn3-cre-tdTomato mice bilaterally injected with AAV-hM4D-mCitrine virus in the MePD to selectively inhibit Ucn3 neurons, with clozapine-N-oxide (CNO) (Tocris Bio-techne, Abingdon, UK), in the presence of acute TMT-exposure or restraint stress. For LH measurement, blood samples were collected every 5 min for 2 h, as described above. CNO was made fresh on the day of experimentation and administered, i.p. in saline at a dose of 5 mg kg^−1^, after 30 min of controlled blood sampling for LH measurement. TMT was introduced into the animal’s cage after 60 min of blood sampling and remained in place for the remaining 1 h of experimentation, as described above. For experiments involving restraint stress, the animals were placed in a restraint device, after CNO injection (at 30 min), 60 min into blood sampling, with restraint and blood sampling continuing for the remaining 1 h duration of the experiment. On a separate occasion, mice received a saline injection at 30 min and were exposed to either TMT or restraint stress, 60 min into the expetriment. For CORT measurments, 15 μl blood samples were collected on a separate occasion using the tail-tip bleeding procedure and blood samples were stored in tubes containing 5 μl heparinised saline (5 IU ml^−1^). CNO or saline were injected at the start of the experiment (time point, −30 min), when the first blood sample was collected. TMT was introduced into the cage, as described above, 30 min into the experiment (time point, 0 min) and lasted for 1 h (time point 60 min). Remaining blood samples were collected at 0, 30 and 60 min, with a final sample taken 1 h after the removal of TMT (time point, 120 min). At the end of the experiment, blood samples were centrifuged at 13,000 RPM for 20 min at 20°C and plasma stored at −20°C. All experiments were performed between 9:00-12:00 h. The mice received CNO or saline in a random order with at least 2 days between experiments for CORT and LH measurment. To control for the effects of the DREADD receptor, the group of Ucn3-cre-tdTomato mice that received the control AAV-YFP, which does not contain the hM4D construct, were administered with CNO in the presence of the stressors and blood samples were collected as described above for CORT and LH measurment. To control for the effects of CNO, control AAV-YFP and DREADD mice were administered CNO alone and blood samples were collected for LH measurement. This study was cross-designed, as described above.

### Validation of AAV injection and cannula implant site

Mice were anaesthetised with a lethal dose of ketamine upon completion of experiments. Transcardial perfusion was performed with heparinised saline for 5 min followed by ice-cold 4% paraformaldehyde (PFA) in phosphate buffer (pH 7.4) for 15 min using a pump (Minipuls, Gilson, Villiers Le Bel, France). Brains were collected immediately, fixed in 15% sucrose in 4% PFA at 4 °C, left to sink and then transferred to 30% sucrose in phosphate-buffered saline (PBS) and left to sink. Brains were snap-frozen in isopropanol on dry ice until further processing. Every third coronal brain section, 30-μm/section, was collected using a cryostat (Bright Instrument Co., Luton, UK) through-out the MePD region corresponding to −1.34 mm to −2.70 mm from bregma. Sections were mounted on microscope slides, air dried and covered with ProLong Antifade mounting medium (Molecular Probes, Inc. OR, USA). Verification of AAV injection site and cannula placement was performed using Axioskop 2 Plus microscope equipped with Axiovision, version 4.7 (Zeiss) by examining whether cannula reach the MePD for neuropharmacological experiments. For AAV-hM4D-mCitrine injected Ucn3-cre-tdTomato mice we determined whether Ucn3 neurons were infected in the MePD region by merging td-Tomato fluorescence of Ucn3 neurons with mCitrine fluorescence in the MePD.

### LH pulse detection and analysis

Processing of blood samples was performed using an LH ELISA as previously reported (32). The capture antibody (monoclonal antibody, anti-bovine LHβ subunit, AB_2665514) was purchased from Department of Animal Science at the University of California, Davis. The mouse LH standard (AFP-5306A) and primary antibody (polyclonal antibody, rabbit LH antiserum, AB_2665533) were obtained from Harbour-UCLA (California, USA). The secondary antibody (Horseradish-Peroxidase (HRP)-linked donkey anti-rabbit IgG polyclonal antibody, AB_772206) was purchased from VWR International (Leicestershire, UK). Intra-assay and inter-assay variations were 4.6% and 10.2%, respectively. The ODs of the standards were plotted against the log of the standard concentrations. Non-linear regression was used to fit the points and various parameters were extracted to calculate the concentration of LH (ng/ml) in the blood samples, as previously described (32). The concentration of LH at every time point of blood collection was plotted as a line and scatter graph using Igor Pro 7, Wavemetrics, Lake Oswego, OR, USA. DynPeak algorithm was used for the detection of LH pulses (33).

### CORT detection

We used a commercially available enzyme immunoassay (EIA) kit (DetectX® Corticosterone Enzyme Immunoassay Kit, K014; Arbor Assays, Michigan, USA) to detect levels of CORT hormone in plasma samples, per the manufacturer’s protocol. The kit was stored at 4°C, upon arrival. A microplate reader (BMG Labtech, Aylesbury, UK) was used to read the optical density of each sample at 450 nm.

### Statistics

For neuropharmacological experiments, female C57Bl6/J mice treated with Ucn3 and Ast2B were compared between groups using a Wilcoxon signed ranks (T) test. For DREADDs experiments, mice administered with saline and CNO were also compared between groups using a Wilcoxon signed ranks (T) test and CNO only data was presented as mean ± SEM. Statistics were performed using Igor Pro 7, Wavemetrics, Lake Oswego, OR, USA. Data was represented as mean ± SEM and +p<0.05, ++p<0.001 and +++p<0.0001 were considered to be significant.

## Results

### Intra-MePD administration of Ucn3 dose-dependently suppresses pulsatile LH secretion in adult OVX mice

Unilateral Ucn3 delivery into the right MePD suppressed pulsatile LH secretion in a dose-dependent manner. On average the mice treated with both doses of Ucn3 exhibited significantly increased LH pulse interval compared to the pre/post-infusion periods (Fig. 1 B, C, and D; 336.60 fmol, ++p<0.001; n=7; 3.36 pmol, +++p<0.0001; n=10). aCSF had no effect on LH pulse interval (Fig. 1, A and D; n=7). These data are summarized in the figure 1D. All cannulae were correctly placed in the MePD.

**Figure 1.**
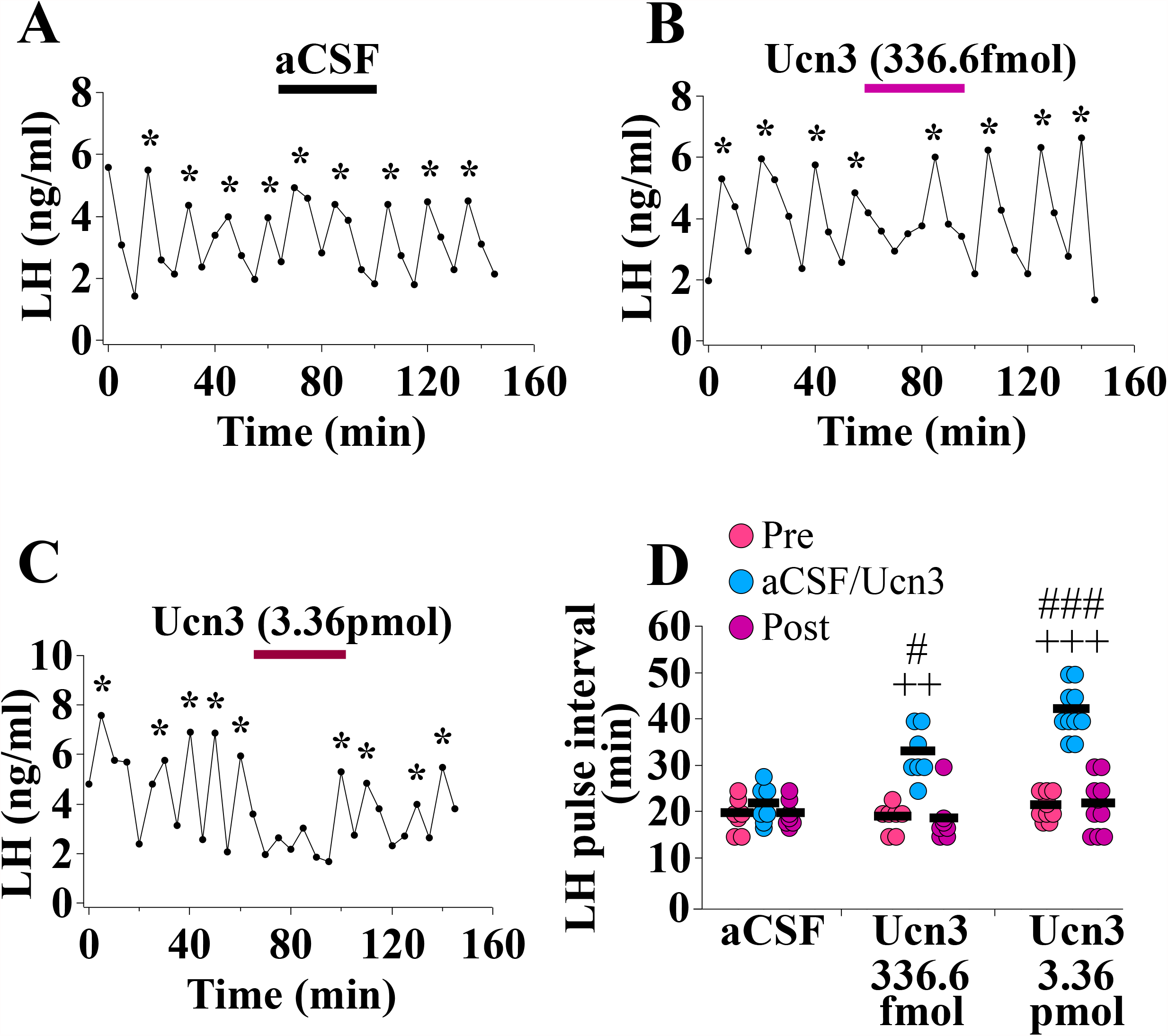
Dose-dependent suppression of pulsatile luteinising hormone (LH) secretion by unilateral intra-posterodorsal medial amygdala (MePD) infusion of Urocortin 3 (Ucn3) in adult ovariectomised (OVX) C57Bl6/J female mice. Representative LH pulse profile with (A) aCSF, (B) 336.60 fmol in 1.80 μl or (C) 3.36 pmol in 1.80 μl of Ucn3 infusion for 35 min administered between 60-95 min of the total blood sampling period. (D), Summary of LH pulse interval for the 60 min pre-infusion control period (0-60 min), 35 minute aCSF/Ucn3 infusion period (60-95 min) and 55 min post-infusion recovery period (95-150 min). LH pulses detected by the DynePeak algorithm are indicated with an asterisk located above each pulse on the representative LH pulse profiles. ++p<0.001, +++p<0.0001 Ucn3 vs pre-treatment control period; #p<0.05, ###p<0.0001 Ucn3 vs aCSF at the same time point; n=7-10 per group.

### Effect of CRF-R2 antagonism on TMT-induced suppression of LH pulsatility

TMT exposure increased LH pulse interval compared to the control period and non-TMT controls (Fig. 2, A, B and E; pre-TMT vs TMT, +++p<0.0001; n=12). Bilateral delivery of Ast2B, a selective CRFR2 antagonist, into the MePD completely blocked the TMT induced suppression of LH pulsatility (Fig. 2, D and E; TMT+Ast2B vs TMT, ###p<0.0001; n=11). Delivery of Ast2B alone had no effect on LH pulsatility (Fig. 2, C and E; n=7), as well as no TMT exposure (Fig. 2, A and E; n=6). The results of this experiment are summarised in the figure 2E. These data show that TMT is probably modulating Ucn3/CRFR2 activity in the MePD to mediate its inhibitory effects on GnRH pulse generator frequency. Mice with misplaced cannulae (n=2) were excluded from the analysis.

**Figure 2.**
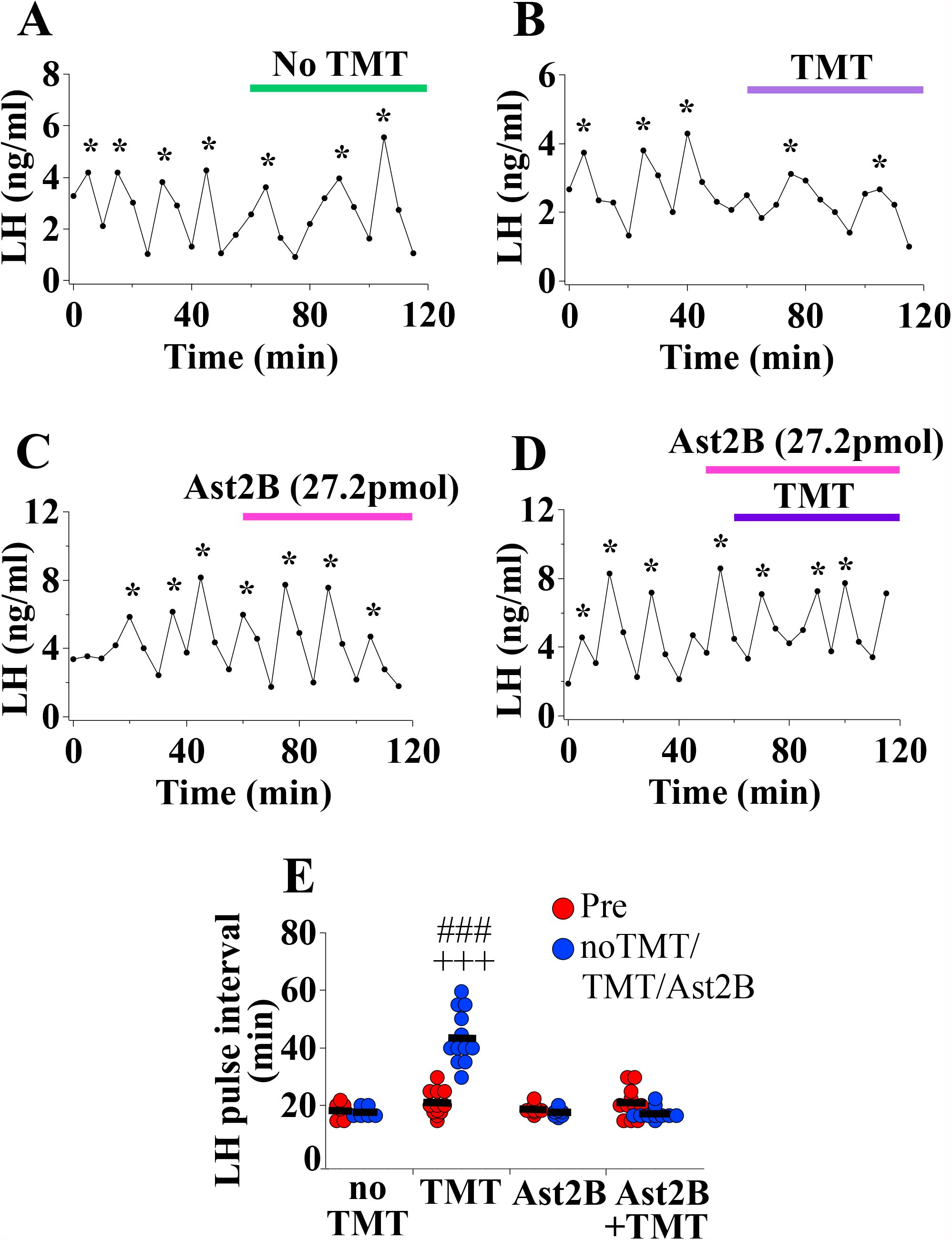
Acute 2,4,5-Trimethylthiazole (TMT)-exposure suppresses pulsatile luteinising hormone (LH) secretion and Astressin2B (Ast2B) delivery into the posterodorsal medial amygdala (MePD) completely blocks the TMT effect on LH pulsatility in adult ovariectomised (OVX) C57Bl6/J female mice. Representative LH pulse profile with (A) control non-TMT (B) TMT-exposure (C) 27.2 pmol in 1.7 μl of Ast2B infusion alone (D) 27.2 pmol in 1.7 μl + TMT Ast2B infusion. (E), Summary of LH pulse interval for the pre-TMT control period (0-50 min), Ast2B infusion (50-120 min) and TMT exposure (60-120min). LH pulses detected by the DynePeak algorithm are indicated with an asterisk located above each pulse on the representative LH pulse profiles. ++p<0.001 pre-TMT vs. TMT; ##p<0.001 TMT vs. no TMT and 16 μM Ast2B; ###p<0.0001 TMT vs TMT+16 μM Ast2B; n=6-12 per group.

### DREADD inhibition of Ucn3 neurons in the MePD blocks the inhibitory effect of TMT exposure on LH pulsatility

Bilateral inhibition of MePD Ucn3 neurons via activation of the hM4D receptor with CNO, in OVX Ucn3-Cre-tdTomato mice, completely blocked the suppressive effect of TMT on pulsatile LH secretion compared to controls (Fig. 3, A, B, C and E; control vs CNO, ##p<0.001; DREADD, n=8, control AAV, n=5, saline, n=5). CNO administration in control AAV-YFP injected mice did not block TMT-induced suppression of LH pulsatility (Fig. 3, C; LH pulse interval, pre-TMT 17.29 ± 1.04 min, TMT 51.25 ± 7.18 min, mean ± SEM). Data for saline and control AAV-YFP injected mice were combined as control since there was no significant difference between the two control groups (Fig. 3, A, C and E, controls pre-TMT vs TMT, ++p<0.001). The data from these experiments are summarised in the figure 3E. Administration of CNO alone had no effect on LH pulsatility (Fig. 3, D; LH pulse interval, pre-CNO 17.50 ± 1.44 min, post-CNO 17.79 ± 1.40 min, mean ± SEM; n=4). The results from this study agree with the neuropharmacological data above showing that the inhibitory effect of TMT on pulsatile LH secretion is likely mediated by Ucn3 activity in the MePD. Mice with misplaced injections (n=2) were excluded from the analysis.

**Figure 3.**
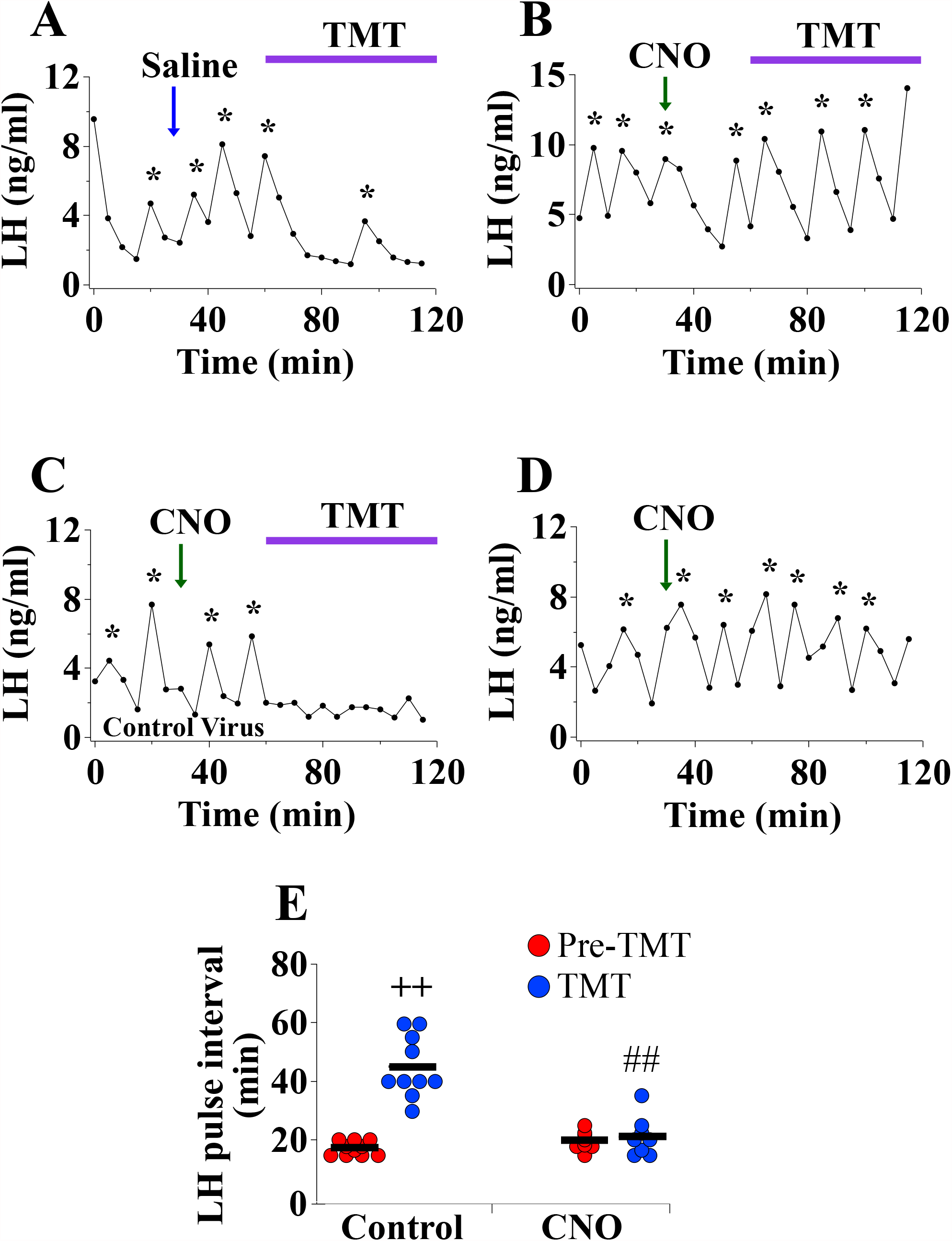
Bilateral DREADD inhibition of posterodorsal medial amygdala (MePD) Urocortin3 (Ucn3) neurons blocks the suppressive effect of 2,4,5-Trimethylthiazole (TMT) on luteinising hormone (LH) pulsatility in adult ovariectomised (OVX) Ucn3-Cre-tdTomato female mice. Representative LH pulse profiles showing the effects of (A) Saline (30 min into the pre-TMT control blood sampling period) and TMT-exposure for AAV-hM4D injected mice, (B) clozapine-N-oxide (CNO) (administered, i.p. in saline at a dose of 5 mg kg^−1^, 30 min into the pre-TMT control blood sampling period) and TMT-exposure (60 min into blood sampling) for AAV-hM4D injected mice, (C) CNO and TMT-exposure for control AAV-YFP injected mice, (D) CNO alone (administered, i.p. in saline at a dose of 5 mg kg^−1^, 30 min into the control blood sampling period) for control AAV-YFP and AAV-hM4D injected mice. E), Summary of LH pulse interval for the pre-TMT control period (1 h) and TMT-exposure period (1 h). LH pulses detected by the DynePeak algorithm are indicated with an asterisk located above each pulse on the representative LH pulse profiles. ++p<0.001 pre-TMT vs TMT for control group; ##p<0.001 control vs. DREADD; n=5-10 per group.

### DREADD inhibition of Ucn3 neurons in the MePD blocks the effect of restraint stress on LH pulsatility

Bilateral DREADD inhibition of MePD Ucn3 neurons in OVX Ucn3-cre-tdTomato mice, completely blocked the inhibitory effect of restraint stress on pulsatile LH secretion compared to saline treated controls (Fig. 4, A, B, C and E; control vs DREADD, ##p<0.001; DREADD, n=7, control AAV, n=5, saline, n=6). CNO administration in control AAV-YFP injected mice did not block TMT-induced suppression of LH pulsatility (Fig. 4, C; pre-restraint 18.67 ± 1.25 min, restraint 58.00 ± 2.55 min, mean ± SEM). Data for saline and control AAV-YFP injected mice were combined as control since there was no significant difference between the two control groups (Fig. 4, A, C and E, control pre-restraint vs restraint, ++p<0.001). The results from these experiments are summarised in the figure 4E. Administration of CNO alone had no effect on LH pulsatility (Fig. 4, D; LH pulse interval, pre-CNO 17.50 ± 1.44 min, post-CNO 17.79 ± 1.40 min, mean ± SEM; n=4). The data from this study shows that Ucn3 activity in the MePD possibly mediates the effect of restraint stress on pulsatile LH secretion.

**Figure 4.**
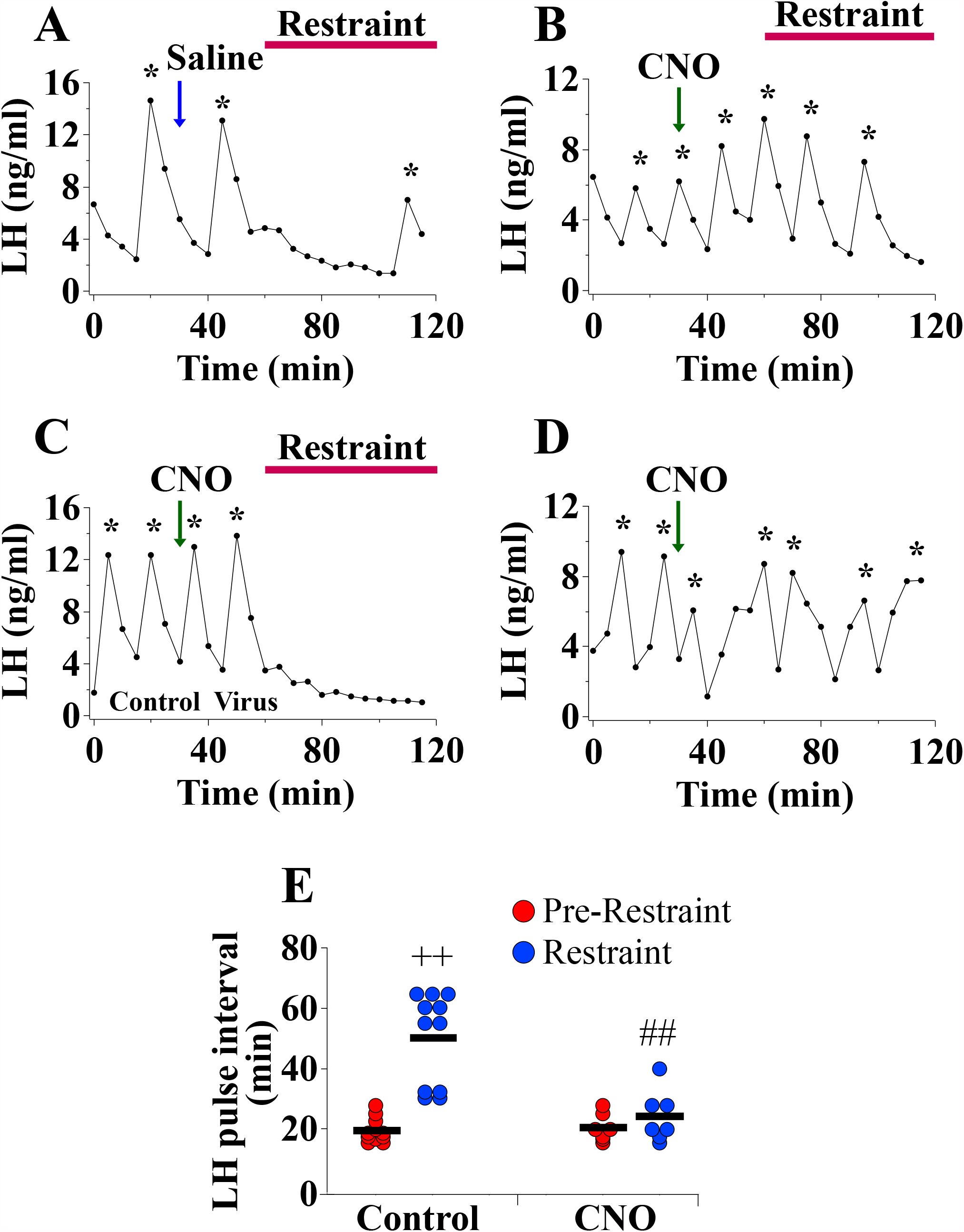
Bilateral DREADD inhibition of Urocortin 3 (Ucn3) neurons in the posterodorsal medial amygdala (MePD) blocks the inhibitory effect of restraint stress on luteinising hormone (LH) pulsatility in adult ovariectomised (OVX) Ucn3-cre-tdTomato female mice. Representative LH pulse profiles showing the effects of (A) saline (30 min into the pre-restraint control blood sampling period) and restraint for AAV-hM4D injected mice, (B) clozapine-N-oxide (CNO) (administered, i.p. in saline at a dose of 5 mg kg^−1^, 30 min into the pre-restraint control blood sampling period) and restraint for AAV-hM4D injected mice, (C) CNO and restraint for control AAV-YFP injected mice, (D) CNO alone (30 min into the control blood sampling period) for control AAV-YFP and AAV-hM4D injected mice. (E), Summary of LH pulse interval for the pre-restraint control period (1 h) and restraint period (1 h). LH pulses detected by the DynePeak algorithm are indicated with an asterisk located above each pulse on the representative LH pulse profiles. **p<0.001 pre-restraint vs. restraint for control group; #p<0.001 control restraint vs. CNO restraint; n=7-11 per group.

### DREADD inhibition of Ucn3 neurons in the MePD blocks the effect of TMT on CORT release

Bilateral DREADD inhibition of MePD Ucn3 neurons in OVX Ucn3-cre-tdTomato mice completely blocked the TMT-induced CORT rise compared to controls at 60 and 120 min measured on a separate occasion to LH pulse sampling (Fig 5, control vs DREADD, **+**p<0.05; DREADD, n=6, saline, n=6, control AAV, n=3). CNO or saline injections were performed at −30 min, TMT exposure was initiated at 0 min and maintained for 1 h. By 30 min there was an observable rise in CORT for the control group. At 60 min CORT was significantly elevated in the control group compared to the DREADD group (Fig 5, control vs DREADD, **+**p<0.05 at 60 min). CORT remained significantly elevated in the saline group compared to the DREADD group at 120 min (Fig 5, control vs DREADD, **+**p<0.05 at 120 min). In mice injected with control AAV-YFP, CNO was not able to block the TMT-induced CORT rise, thus data for saline and control AAV-YFP injected mice were combined together as control, as there was no significant difference between the two control groups (Fig 5, control). These data show that Ucn3 activity in the MePD mediates the effect of TMT on CORT release.

**Figure 5.**
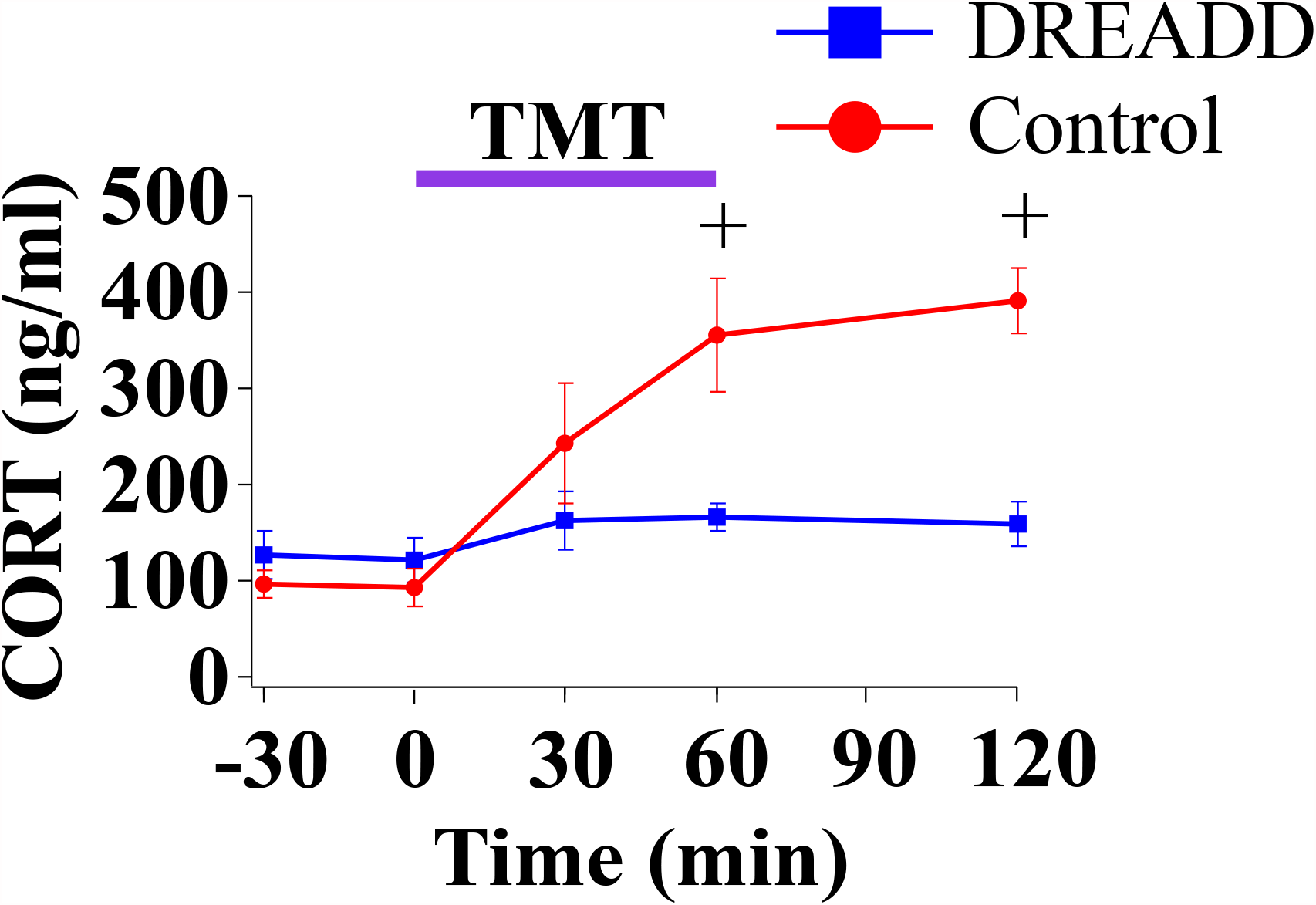
Bilateral DREADD inhibition of Urocortin 3 (Ucn3) neurons in the posterodorsal medial amygdala (MePD) blocks the 2,4,5-Trimethylthiazole (TMT)-induced rise in corticosterone (CORT) in adult ovariectomised (OVX) Ucn3-cre-tdTomato female mice. CORT secretion time-course for mice injected with clozapine-N-oxide (CNO), administered, i.p. in saline at a dose of 5 mg kg^−1^, at the start of the experiment and exposed to TMT, lasting for 1 h, followed by a 1 h recovery period (60-120 min) for the DREADD group (blue line, squares) and control group (red line, circles). CNO or saline were administered at −30 min. TMT-exposure was initiated at 0 min and terminated at 60 min. +p<0.05 control vs. DREADD group at 60 min and 120 min; n=6-9 per group.

### Selective expression of DREAD(Gi) in MePD Ucn3 neurons

Evaluation of m-Citrine, hM4D, expression in tdTomato labelled neurons from AAV-injected Ucn3-cre-tdTomato mice revealed that 83 ± 5% of MePD Ucn3 neurons expressed hM4D (Fig 6, A-F, n=8).

**Figure 6.**
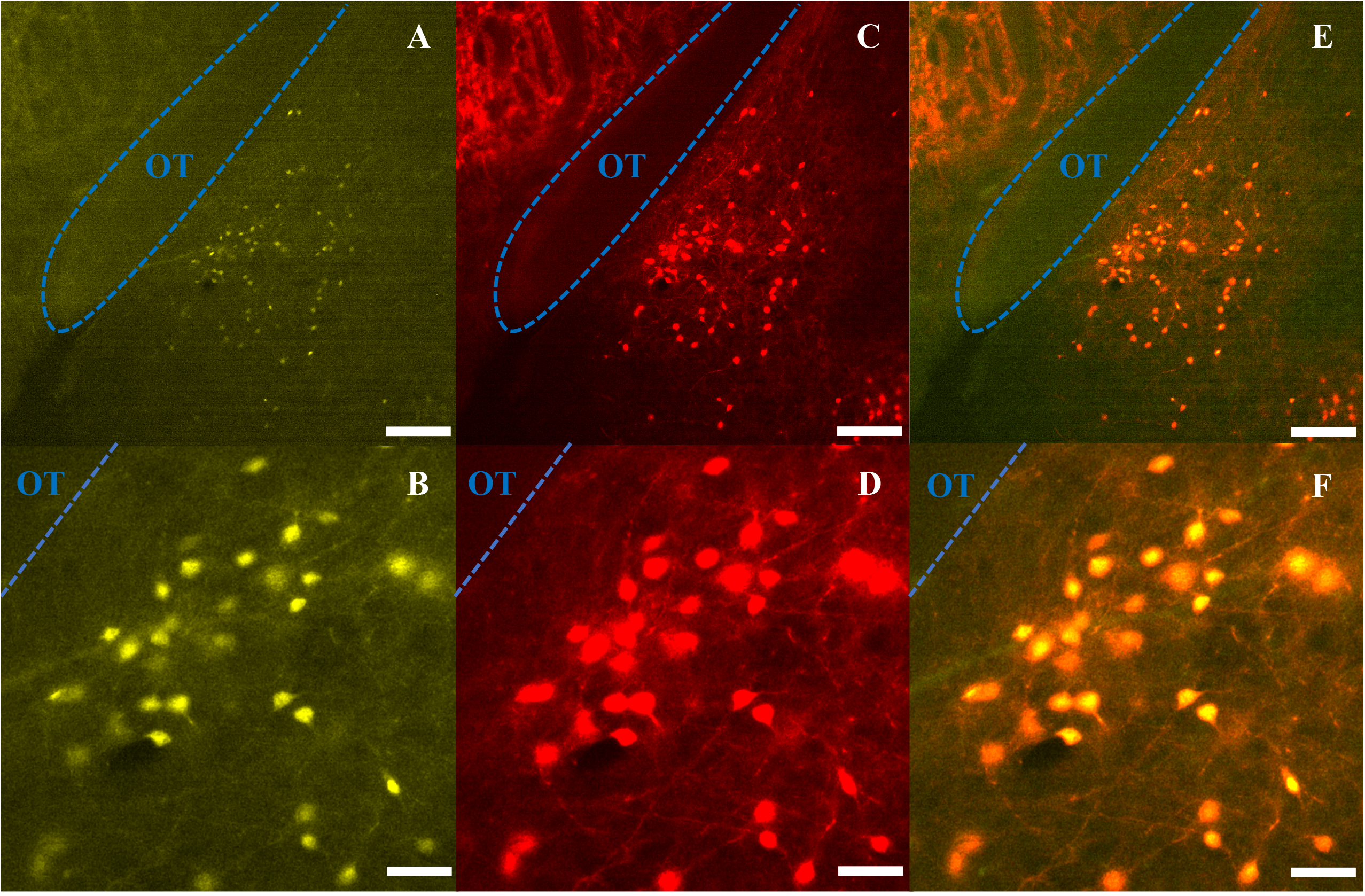
Expression of AAV-hM4D(Gi)-mCitrine in posterodorsal medial amygdala (MePD) Urocortin3 (Ucn3) neurons. (A-F) Representative dual fluorescence photomicrographs of the MePD from a Ucn3-cre-tdTomato female OVX mouse injected with AAV5-hSyn-DIO-HA-hM4D(Gi)-IRES-mCitrine. Ucn3 neurons labelled with m-Citrine (A) and tdTomato (C) appear yellow/orange (E). (B), (D) and (F) are a higher power view of (A), (C) and (E) respecitively. Scale bars represent A, C, E 100 μm and B, D, F 25 μm; OT, Optic tract (blue line).

## Discussion

In the present study, we show intra-MePD administration of the stress neuropeptide Ucn3 dose-dependently suppresses GnRH pulse generator frequency. Additionally, acute exposure to predator odor decreased pulsatile LH secretion, an effect that was completely blocked by CRFR2 antagonism with Ast2B in the MePD. Moreover, DREADD inhibition of Ucn3 neurons in the MePD blocks acute predator odor and restraint stress-induced inhibition of LH pulses and the rise in CORT secretion. We demonstrate for the first time that acute exposure to psychological stressors, which robustly suppresses the GnRH pulse generator is mediated by Ucn3-CRFR2 neural circuitry in the MePD of OVX mice.

Stress exerts a profound inhibitory effect on reproductive function. The amygdala coordinates stress, anxiety and fear states, which regulate pulsatile reproductive hormone release modulating the fertility of species in adverse environments (34). We have previously shown that the MePD is a major regulator of sexual motivation and anxiety behaviour (35) as well as pubertal timing (5,36). Electrophysiological recordings reveal the MePD is activated in response to predator urine (23) or restraint stress (37), which is associated with suppressing LH secretion (4) and LH pulsatility (2). Psychosocial stress decreases c-fos expression in ARC kiss1 neurons, which is linked to a reduction of pulsatile LH secretion (10). Our aim was to elucidate the neural circuitry within the MePD involved in processing stress and relaying this information to suppress the hypothalamic GnRH pulse generator.

Ucn3 and CRFR2 neuronal populations have been identified within the MePD (27). Restraint stress increases Ucn3 mRNA expression (28) and c-fos in the MePD (38) and predator odor activates CRFR2 positive cells in the MeA of rodents (29). In the present study we found that intra-MePD administration of Ucn3 dose-dependently suppresses pulsatile LH secretion and CRFR2 antagonism in the MePD blocks the inhibitory effect of TMT on LH pulsatility. This is consistent with previous studies showing the involvement of CRFR2 signalling in mediating stress signals to the GnRH pulse generator, since central antagonism of CRFR2, using an intracerebroventricular route, blocks the inhibitory effect of stress on LH pulses in rodents (39). Moreover, *in-vitro* application of CRFR2 agonists decreases GnRH neuronal firing rate (40). We show that inhibiting Ucn3 neurons in the MePD prevents the suppressive effect of acute TMT or restraint stress exposure on pulsatile LH secretion. These findings reveal a novel role for MePD Ucn3 neurons as mediators of diverse psychological stress signals modulating GnRH pulse generator activity.

The MePD is of pallidal embryological origin and has a major GABAergic output to key hypothalamic reproductive nuclei (14,41), which might include the ARC (3). Moreover, stress decreases inhibitory GABAergic control within the amygdala leading to enhanced inhibitory efferent output from the amygdala (42,43). Kisspeptin signalling in the MePD regulates GnRH pulse generator frequency (44) and recently we have shown selective optogenetic stimulation of MePD kiss1 neurons increases LH pulse frequency (19). Direct neural projections, of unknown phenotype, extend from the MePD to kiss1 neurons in the ARC (17,18). This connection to the KNDy network might provide a route by which the MePD integrates incoming anxiogenic signals with GnRH pulse generator function. Therefore, we propose that modulation of the reproductive axis by the MePD may involve a disinhibitory system whereby kiss1 neuron activity in the MePD stimulates inhibitory GABAergic interneurons which in turn inhibit GABAergic efferents from the MePD to the ARC, thus allowing for the observed increase in GnRH pulse generator frequency in response to activation of MePD kiss1 neurons (19). Conversly, reduced kisspeptin signalling in the MePD, as with kisspeptin antagonism, decreases GnRH pulse generator frequency (44).

Within the MeA, local Ucn3 fibres overlap with sites expressing CRFR2 (45,46) and MeA Ucn3 neurons have been shown to include an interneuron-like appearance (26) indicating that Ucn3 neurons may connect to and signal via CRFR2 neurons in the MeA. Our neuropharmacological observation and chemogenetic data where antagonism of CRFR2 and inhibition of Ucn3 in the MePD block the effect of TMT on LH pulsatility suggests the activity of both Ucn3 and CRFR2 within the MePD is necessary for mediating the suppressive effects of stress on GnRH pulse generator activity. We know the majority of CRFR2 neurons in the MePD co-express GAD65 and 67 (27), which are required for the synthesis of GABA indicating MePD CRFR2 neurons are likely GABAergic. Based on our findings we propose that stress activated Ucn3 neurons may signal through inhibitory GABAergic CRFR2 neurons upstream to kiss1 neurons in the MePD, thus downregulating kiss1 signalling which in turn decreases GnRH pulse generator frequency. However, we cannot exclude the possibility of alternative routes via which the Ucn3 and CRFR2 neurons in the MePD may modulate GnRH pulse generator activity.

We have shown for the first time that TMT increases and maintaines elevated CORT secretion in mice. The TMT-induced rise in CORT was maintained for at least 1-hour after termination of the predator odor stress, indicating prolonged HPA axis activation, which has been observed with prolonged restraint stress in mice (47). This indicates exposure to these psychogenic stressors can lead to prolonged HPA axis activation in mice. CRFR2 knock-out (KO) mice show an early termination of adrenocorticotropic hormone (ACTH) release in reponse to restraint stress exposure suggesting CRFR2 signalling is important for maintaining HPA axis drive (48). This may provide a potential mechanism by which CRFR2 mediates the prolonged HPA axis activation we observed in response to predator odor. Contrastingly, a commonly held view is that CRFR2 signalling underlies stress-coping mechanisms to counter the stress provoking effects of CRFR1. However, many earlier observsations were based on experimental manipulations lacking absolute region and receptor specificity, which are complicated by compensatory or pleiotropic effects, but the rise in site-specific manipulation of the CRFR1 and CRFR2 receptor signalling in recent reports contradicts the traditional view (49). Furthermore, we show that inhibition of Ucn3 neurons in the MePD not only blocks the suppressive effect of TMT on LH pulsatility, but also prevents the rise in CORT secretion induced by TMT exposure. These data show that Ucn3 signalling in the MePD plays a dual role in modulating GnRH pulse generator activity while interacting with the HPA axis to regulate CORT secretion in response to stress, revealing the MePD as a tenable central hub for integrating incoming anxiogenic signals with the reproductive and stress axes. Our physiological observations will provide valuable parameterisation for our recent mathematical model integrating these neuroendocrine axes (50), adding the MePD Ucn3-CRFR2 system to the network architecture to provide mechanistic details underlying the dynamic relationship between CORT levels and suppression of GnRH pulse generator activity.

Central administration of Ucn3 has been shown to increase CRF and vasopressin concentration in the hypothalamus and elevate CORT secretion in mice (51), and augment the HPA axis response to restraint stress in rats (28). TMT increases c-fos mRNA expression in the MeA (52) and restraint stress is shown to increase specifically Ucn3 mRNA expression in the MeA (28). The MeA projects to the paraventricular nucleus (PVN) of the hypothalamus to activate the HPA axis in response to predator odor (22) and lesioning of the MeA eliminates defensive responses to predator odor in rodents (53). Retrograde tracing has also revealed that the MePD sends stress-activated efferents to the PVN (38). More recently, Ucn3 and CRFR2 positive neurons in the MePD have been shown to receive inputs from and project to the PVN (27), providing a possible direct route for the interaction of MePD Ucn3 with the HPA axis. We show the Ucn3 neuronal population in the MePD is involved in activating the HPA axis in response to predation threat, uncovering a novel functional role for MePD Ucn3 neurons in modulation of the HPA axis during psychosocial stress exposure.

Our findings show for the first time that Ucn3-CRFR2 signalling in the MePD mediates the suppressive effect of psychosocial stress on GnRH pulse generator frequency as well as activating the HPA axis. How the Ucn3 and CRFR2 positive neuronal populations in the MePD communicate with the GnRH pulse generator to suppress pulsatile LH secretion and crosstalk with the HPA axis to modulate CORT in response to psychosocial stress remains to be established. Investigating Ucn3 and CRFR2 stress signalling in the limbic brain has furthered our understanding of the neural mechanisms by which stress exerts its inhibitory effects on the reproductive system, providing an important contribution to the field of reproductive nueroendocrinology that may ultimately improve our treatment options for stress-induced infertility.

## Acknowledgements

The authors gratefully acknowledge the financial support from UKRI: BBSRC (BB/S000550/1) and MRC (MR/N022637/1). DI is a PhD student funded by a MRC-DTP studentship at King’s College London.

